# Optimization of H9c2 differentiation leads to calcium-active and striated cardiac cells without addition of retinoic acid

**DOI:** 10.1101/2024.09.24.614681

**Authors:** Judith Brock, Marcel Hörning

## Abstract

As a reliable alternative to animal testing in cardiovascular research, it is crucial to improve differentiation of immortalized cell lines. In this study, we focused on optimizing the differentiation efficiency of the H9c2 cell line into cardiomyocytes using a high-throughput, automated image processing approach. While previous studies used protocols involving retinoic acid to enhance cardiac differentiation, we applied a simplified medium composition that results in higher differentiation rates. Along that line, we differentiated H9c2 cells into cardiomyocytes, which not only showed sarcomere-characteristic striation but also periodic intracellular calcium signaling for the first time. As a second step, we examined the potential application of polyacrylamide hydrogels (E = 12 kPa) with defined fibronectin coating densities. The optimum fibronectin density of 2.6 µg/cm^2^ found for single cells was investigated to further improve the differentiation efficiency. However, the differentiation and proliferation dynamics dominate the adhesion forces between the cells and the hydrogel, and thus, result in premature clustering and detachment. In conclusion, we identified an optimized differentiation protocol and provided a basis for the further investigation necessary to potentially use hydrogels as natural cell environment, aiming to raise the differentiation efficiency even more.

## Introduction

Cardiovascular diseases have been the leading cause of death worldwide for decades. The necessity to find and improve methods of treatment and to conduct research in the field of cardiac regenerative medicine is thus immense. Typical functional cardiac diseases are hypertrophy, arrhythmias and cardiomyopathies (Ravi et al., 2021). Primary heart cells, particularly cardiomyocytes, are often used to gain knowledge about fundamental heart conditions (Nakayama et al., 2007; Giguère et al., 2018; Hörning et al., 2012). As alternative cell systems, biomedical studies regularly investigate embryonic stem cells (ESCs) and induced pluripotent stem cells (iPSCs), which can differentiate into distinct cell types including cardiac cells (Boheler et al., 2002; Ohno et al., 2013; Narazaki et al., 2008). While both primary and stem cells exhibit similar morphology and behavior as *in-vivo* cells, they also have downsides like ethical concerns, tumor formation, or high expenses (Peter et al., 2016; Nori et al., 2015). Therefore, immortalized cardiac cell lines, such as HL-1, AC16, or H9c2, still serve as essential tools in research (White et al., 2004; Davidson et al., 2005; Kimes and Brandt, 1976).

H9c2 cells were originally isolated from embryonic rat ventricular tissue and are of special interest because they can differentiate into both skeletal and cardiac muscle cells (Kimes and Brandt, 1976; Ménard et al., 1999; Branco et al., 2015). The differentiation into cardiomyocytes is triggered by the addition of retinoic acid to medium with a low serum concentration. So far, this cell line has been used to investigate cardiotoxicity (Daubney et al., 2015; Witek et al., 2016), cardiac hypertrophy, (Watkins et al., 2011; Xu et al., 2020), oxidative stress (Pesant et al., 2006), and calcium channels (Ménard et al., 1999; Hescheler et al., 1991). However, even though these cells express cardiogenic markers like alpha actinin, myosin light chain 2, and cardiac Troponin T, they lack the formation of sarcomeric structures and do not show contractility (Suhaeri et al., 2015). Additionally, differentiation of these cells is complex, and the H9c2 cell line might even resist differentiation, as previously suggested (Patten et al., 2017). Finding supplementary or optimized ways for differentiation is therefore necessary to use this cell line more reliably as a model for cardiomyocytes and to potentially learn more about heart diseases.

For the optimization of H9c2 differentiation the extracellular matrix (ECM) is one starting point, which has not been sufficiently investigated. However, numerous studies have identified the crucial influence between cellular dynamics and the properties of the ECM Lukashev and Werb (1998); Reilly and Engler (2010); Chaudhuri et al. (2020). The ECM serves as a three-dimensional scaffold which initiates mechanical and biochemical cues cells need during adhesion, proliferation, migration, and differentiation (Frantz et al., 2010; Lukashev and Werb, 1998; Chaudhuri et al., 2020). During all of these cellular dynamics an interplay between matrix properties, i.e., elasticity, topography, and several proteins, takes place. Regarding cell differentiation, the rigidity of the ECM guides the cells, evoking a higher differentiation efficiency on substrates with tissue-like stiffness (Engler et al., 2004). Mesenchymal stem cells, for example, differentiate into varying cell types in connection to the stiffness of the substrate. On soft hydrogels they form neurons, on hydrogels with medium rigidity, they transform to myocytes and on hard substrates to osteocytes (Smith et al., 2018; Engler et al., 2006). As cells are sensitive to their surroundings, they can likewise detect the topography of the ECM and are able to orient and align themselves in accordance to micro and nanostructures like patterns, pillars, grooves, and channels (Hume et al., 2012; Connon and Gouveia, 2021; Yamamoto et al., 2008). These topographical structures can be applied either by roughening the surface of the substrate or by applying a predesigned pattern and aim to mimic the natural environment cells rely on during differentiation (Cui et al., 2021). In this connection, an advanced myogenic maturation and sarcomere formation compared to control cells plated on unpatterned gels has been exemplified for myoblasts plated on micro patterned hydrogels (Denes et al., 2019; Bettadapur et al., 2016; Engler et al., 2004). Similarly, the presence of fibrous ECM proteins, like laminin, elastin, collagen, and fibronectin, plays a vital role in differentiation by enhancing the amount of cells that turn into a desired cell type and by affecting the switch between proliferation and differentiation (García et al., 1999; Li et al.). C2C12 cells, for instance, display a relation between the the fibronectin concentration and the amount of myogenic differentiation markers (Salmerón-Sánchez et al., 2011) and adhesion area (Brock et al., 2022).

The aim of this study is to improve H9c2 differentiation efficiency. For that we first investigated cells on glass focusing on the culture conditions, such as medium exchange and composition, to ensure the comparability to former studies. Applying high-throughput, automated image processing analysis, we found that normal growth medium with 10% FBS led to the highest amount of cells that differentiated into cardiomyocytes after 2 weeks. In contrast, medium with different RA concentration failed to induce differentiation in a similar manner. The cardiac phenotype was confirmed by an independent test using of the cardiac Troponin T marker. Additionally, we observed muscle-characteristic striation with the sarcomeric alpha actinin marker, and an intracelluar calcium response to extracellular, electrical stimulation. Using the best culture conditions found on glass substrates, we further examined influence of polyacrylamide hydrogels (*E* = 12 kPa) and the fibronectin density on single cells. An optimum was identified at about 2.6 µg/cm^2^ similar to C2C12 cells before (Brock et al., 2022). Combining the optimum culture and fibronectin density conditions, we then compare the differentiation efficiency between glass and hydrogel. Although an improvement was expected, fewer cells differentiated on the optimized ECM.

Nevertheless, the results of this study signify important progress for differentiation of H9c2 cells and can be used as basis for further investigation in more natural cell environments.

## Material and Methods

### Glass Preparation

Round and square-shaped glass cover slips (∅ 22mm, Carl Roth, P235.1; 24 × 24 mm^2^, VWR, ECN632-1571) were cleaned following a modified RCA method (Kern & Puotinen, 1970). Briefly, glasses were successively sonicated in acetone, ethanol, methanol and distilled water for 3 minutes each, then covered with a hydrogen peroxide solution (H2O:H2O2:NH3 aq. as 5:1:1), sonicated for 3 min, followed by incubation at 60°C for 30 min, and finally rinsed 10 times with distilled water and dried completely at 70°C. Round glass substrates were subsequently prepared to bind the hydrogels by incubation in 5% (V/V) vinyltrimethoxysilane (Sigma, 235768) in toluene on a shaker for 18h in the dark at RT. Afterwards the round glasses were washed consecutively with acetone, ethanol and distilled water before being dried at 140°C for at least 1h (Hörning et al., 2017).

### Hydrogel Preparation

For the polyacrylamide hydrogels, a 2% bisacrylamide solution (bAAm, Carl Roth, 3039.2) was added to a 40% acrylamide solution (AAm, Carl Roth, 7748.1) in distilled water at a crosslinker ratio of 2%. The polymerization of this solution was initiated by 10% ammonium persulfate (APS, Sigma, A3678) and N,N,N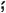N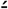tetramethyl ethylenediamine (TEMED, Carl Roth, 2367.3). For the plane hydrogels, 20 µL of this polymerizing solution was then sandwiched between a vinyl-silanized glass cover slip (∅ 22mm) and an RCA cleaned cover slip (24 × 24 mm^2^) and left to polymerize for 30 min at RT. For the mechanically-roughened hydrogels the same protocol was used, except that the square cover slip was mechanically roughened before the application of 40 µL of the polymerizing solution. After polymerization, the hydrogels were soaked in water for at least 48 h to remove residual chemicals. The hydrogels were about 60 µm (plane) and 100 µm (mechanically roughened) thick, as confirmed by microscopy. The elastic modulus was set to about 12 kPa by the concentrations of acrylamide and bisacrylamide solutions (Brock et al., 2022).

### Mechanical Testing

The E-modulus of the hydrogels were measured by nanoindentation using an atomic force microscope (AFM, NanoWizard, JPK Instruments, Berlin, Germany). A silicon nitride cantilever with an attached spherical colloidal probe (CP-PNP-BSG; 0.08 N/m; R = 5 µm, Olympus Optical) was used. The spring constant of the cantilevers was obtained through thermal noise measurements. The indentation curves were measured with an approach speed of 1 µm/s. The data were analyzed using nonlinear least-squares fitting to the Hertz model (Sneddon, 1965; Domke and Radmacher, 1998) with a customized MATLAB (2023b, The MathWorks, Natick, MA, USA) routine. Statistical significance was ensured by the quantification of the Young’s modulus E at 50 independent indentation sites in two 100 × 100 µm^2^ areas for each hydrogel (Brock et al., 2022; Erhardt et al., 2024).

### Surface Functionalization

The hydrogels were functionalized by 3,4-Dihydroxy-L-phenylalanine (L-DOPA, Sigma-Aldrich, D9628) Brock et al. (2022). Briefly, L-DOPA was dissolved in freshly prepared TRIS buffer (10 mM, pH 10, Roth 4855.2) at a concentration of 2 mg/mL for 30 min in the dark on a tube roller and sterilized through a 0.2 µm filter (Filtropur S0.2, Sarstedt 83.1826.001). Followed by a washing with TRIS buffer, 250 µL of this solution was added onto each hydrogel and incubated for 30 min in the dark at RT. To remove unbound L-DOPA, the samples were washed twice with PBS and finally incubated with human plasma fibronectin (Sigma, F2006) at different concentrations for at least 2 h at 37°C.

### Cell Culture

Rat cardiomyoblasts (H9c2 (2-1), <15 passages, Sigma-Aldrich, 88092904) were maintained in Dulbecco’s Modified Eagle’s Medium (DMEM) low glucose (Gibco, 31885023), supplemented with 10% fetal bovine serum (FBS, Gibco, 1027010) and 1% penicillin-streptomycin (Gibco, 15140122) at 37°C and 5% CO_2_ in a humidified atmosphere. Cells were kept at 60-70% confluency and passaged every 2-3 days to retain differentiation potential. For single cell experiments, cells were plated at cell densities between 5 and 13 cells/mm^2^ on 12 kPa substrates coated with different fibronectin densities (*ρ*_FN_ = 0.4 − 4.0 µg/cm^2^) and incubated for 24h before fixation. For differentiation, cells were plated at a cell density of 100 cells/mm^2^ on fibronectin-coated glass substrates (0.4 µg/cm^2^) and hydrogels (2.6 µg/cm^2^) and left to reach confluence in DMEM containing 10% FBS for 3 days. Then the medium was either left at 10% FBS for 14 additional days or switched to DMEM containing 1% FBS with or without the addition of different concentrations (10 - 1000 nM) of all-trans-retinoic acid (RA, Sigma, R2625). This 1% FBS medium was either left for 14 additional days or changed and supplemented with RA every 2-3 days for 2 weeks in total. RA was prepared in DMSO as a 10 nM stock, diluted before each use in DMEM with 1% FBS and added in the dark to prevent degradation.

### Cardiomyocyte Isolation

The primary heart tissues were prepared following a method described in (Hörning et al., 2017, 2012; Erhardt et al., 2024). Briefly, hearts of 1-3-day-old Wistar rats were isolated and consecutively cleaned, minced, and enzymatically digested in five cycles using collagenase type I (Gibco, 17100017). The isolated cells from the last four cycles were pre-plated for 1 h in plastic dishes to reduce the fraction of fibroblasts. Cells were plated at a density of 2.6 × 10^5^ cells/cm^2^ in Dulbecco’s modified Eagle’s medium (DMEM, Gibco, 11885084) supplemented with 10% fetal bovine serum (Gibco, 10270106), 1% penicillin-streptomycin (Gibco, 15140122) and kanamycin sulfate (80 mg/L; Gibco, 11815024). After 24 h, the medium was exchanged to minimal essential medium (MEM, Gibco, 11095080) supplemented with 10% calf serum (Gibco, 16170-087), 1% penicillin-streptomycin, kanamycin sulfate (80 mg/L) and cytosine arabinofuranoside (ARA-C, 10 µM; Sigma-Aldrich, C1768). After 5 days of incubation, the cardiac tissues were fixed and stained, following the same protocol used for differentiated H9c2 cells.

### Fluorescence Staining

Cells were washed with PBS and fixed with 4% formaldehyde in PBS (Thermo Scientific Chemicals, J60401.AK) for 10-20 minutes at RT. For F-actin staining, cells were then washed 3 times with 0.1% Tween 20 (Carl Roth, 9127.1) in PBS for 10 minutes each, labeled with rhodamine phalloidin (0.25 U/mL in methanol, Alexa fluor 546, Invitrogen, A22283) and DAPI (1 µg/mL in PBST, Sigma D9542) for 1h at RT in the dark, subsequently washed 3 times with 0.1% PBST for 10 min, and covered with ProLong Gold antifade reagent (Invitrogen, P10144) until observation. For antibody staining, after fixation, the samples were blocked with 400 µL 0.1% BSA (Sigma Aldrich, A9418) in 0.1% Saponin (Sigma, S1252) for 30 min, then incubated with 200 µL mouse monoclonal anti-sarcomeric alpha actinin (1:200; Invitrogen, MA1-22863) or cardiac troponin T monoclonal antibody (1:200; Invitrogen, MA5-12960) in 0.1% BSA in 0.1% Saponin for 1h at RT, followed by 200 µL secondary antibody, Alexa Fluor 488 goat anti-mouse IgG (1:200; Invitrogen, A11001), rhodamine phalloidin, and DAPI in 0.1% BSA in 0.1% Saponin for 1h at RT. ProLong Gold antifade reagent (Invitrogen, P10144) was used to preserve the samples until observation.

### Image Acquisition of Fluorescent Cells

Fluorescence was observed by an AxioObserver SD confocal microscope (Carl Zeiss Microscopy GmbH Jena, Germany) equipped with a Yokogawa CSU-X1 spinning disk unit at ×40 (Plan-Apochromat 1.4 Oil DIC UV, Zeiss) and using an Axiocam 503 Mono CCD camera (Zeiss) at a resolution of Δ*x* = Δ*y* = 0.227 µm.

The images were acquired and analyzed by the ZEN blue v2.3 software (Zeiss). The cell structures were visualized using the 405 nm (DAPI), 488 nm (Alexa fluor 488) and 561 nm (Phalloidin) diode lasers. The images were obtained as a 5 × 5 tile composition with 11 - 13 focal heights (Δ*z* = 1 µm). The tiles were stitched and then orthogonally projected by the “fuse tiles”, “correct shading” and “orthogonal projection” features of the ZEN 2.3 software.

### Observation of Calcium Transients

For the observation of calcium transients, the differentiated H9c2 cells were incubated with 200 µL of the fluorescence dye Fluo8 (8.3 µM in PBS, AAT Bioquest) at room temperature (RT) in the dark for 30-60 min. Then, the observation was conducted at RT in Tyrode’s solution (136.9 mM NaCl, 1 mM MgCl2, 2.7 mM KCl, 1.8 mM CaCl2, 0.4 mM NaH2PO4 and 5.5 mM glucose (Sigma-Aldrich, T2145)) with additional 2.7 mM KCl (final concentration of 5.4 mM) and 5 mM HEPES (Roth, 9105.2). The pH level was adjusted to 7.4 using NaOH. The calcium transients were observed by a customized microscope setup (ThorLabs) equipped with a Kinetix sCMOS high-speed camera (Photometrics, 140 FPS and Δ*x* = Δ*y* = 48 µm/px after 4 × 4 binning) and a zoom objective (PlanApoZ 0.5 ×/0.125 FWD 114 mm, Carl Zeiss). The samples were electrically stimulated using platinum electrodes (∅ 0.5 mm, 99.997%, Thermo Fischer Scientific/ Alfa Aesar) and a modified version of the MyoPulser stimulator introduced by Ott and Jung (2023). The device was constructed using a motor controller (5 − 35 V, *I*_max_ = 2 A) and a microcontroller (ESP32-S2-WROVER, Espressif) programmed with an Arduino IDE (2.3.2). A power supply unit with a voltage of 12 V and current up to 2 A was used. A customized Graphical User Interface (GUI) was implemented using the software Processing (The Processing Foundation). The device was enclosed in a customized 3D-printed chassis (PRUSA MK3s, Prusa Research), which was designed using the CAD software SolidWorks (Dassault Systemes) (Erhardt et al., 2024). The tissues were paced with monophasic pulses of 10 ms or 50 ms and amplitudes of ± 10 V. The electrodes were approximately 1 cm apart, resulting in an electrical field of 10 V/cm.

### Actin Quantification Analysis (AQuA)

The quantification of actin filaments in muscle cells using AQuA has been described before using customized routines in MATLAB (2023b, The MathWorks, Natick, MA, USA) (Zemel et al., 2010; Inoue et al., 2015; Hörning et al., 2017; Erben et al., 2020; Brock et al., 2022). Briefly, the Laplacian filter

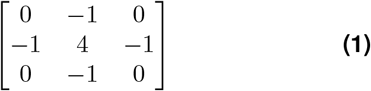

and *n* = 15 differently rotated anisotropic Gaussians with *σ*_*x*_ = 2 px and *σ*_*y*_ = 6 px were convoluted to elongated Laplace of Gaussian (eLoG) kernels. The kernels were applied to the original 5 × 5 tiled images and the maximum response of each pixel was calculated, as

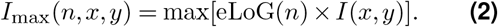

Thereafter, *I*_max_ was processed by the binarized original images using the Otsu’s thresholding method Otsu (1979). Connected fibers of the same rotational direction with less than 10 px for single cells and 20 px for tissues were removed. The obtained actin fibers were colorized with a color scheme that corresponds to the local actin orientation angles, *θ*_*n*_.

### Calcium Transient Analysis

The recorded movies were pre-processed by background subtraction, averaged in time (10 frames), and filtered in space with Gaussian blur (10 px) using a customized macro in ImageJ (1.54f) (Loppini et al., 2022; Erhardt et al., 2024). Further analysis was performed using customized routines in MATLAB (2023b, The MathWorks, Natick, MA, USA). The calcium transients were analyzed at a normalized calcium intensity of 50% of each individual wave to obtain the individual calcium transient duration (CTD) and calcium transient intervals (CTI). From the CTDs and CTIs, the mean values and standard deviation was calculated for each calcium transient.

The normalized calcium transients were calculated by the improved signal oversampling analysis (Erhardt et al., 2024). The periodic signals were pixelwise-stacked by equidistant time intervals, and the CTD was computed from the stacked period-2 calcium transient, as

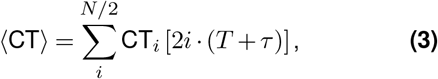

where CT_*i*_ are the calcium transients of the individual waves, *T* is the pacing period, and *τ* is the offset induced by frame rate inaccuracy of the camera (*τ* ≤ 140 Hz ≃ 7 ms).

### Fourier Transformation Imaging (FFI)

FFI was applied to the raw unprocessed fluorescence recordings, as introduced before (Hörning et al., 2017; Loppini et al., 2022; Erhardt et al., 2024). Briefly, the images were decomposed pixelwise and transformed to the mathematically complex Fourier space, *F*_x,y_, as a function of the frequency *f*, i.e., *I*_x,y_(*t*) → *F*_x,y_(*f*), with the intensity *I*_x,y_(*t*) at the spatial position (*x, y*). From that, the amplitudes were calculated and spatially re-composed to a Fourier frequency-series. The amplitude map was selected at the respective frequency *f*_*T*_ = *T* ^−1^ with the pacing period *T*, as |*F* (*f*_*T*_)|.

## Results

The H9c2 rat cardiomyoblast cell line presents a special challenge for differentiation since it can form both cardiac and skeletal muscle cells. In this connection, several studies described that both a reduction of fetal bovine serum (FBS) to 1% and supplementation with retinoic acid (RA) is necessary to obtain cardiomyocytes instead of skeletal muscle cells (Ménard et al., 1999; Branco et al., 2012; Pereira et al., 2011). However, the added RA concentrations range anywhere between 10 and 1000 nM, and the frequency of medium and RA exchange varies from study to study (Ménard et al., 1999; Pereira et al., 2011; Suhaeri et al., 2015; Campero-Basaldua et al., 2023). Therefore, it is still unclear which condition leads to the optimal differentiation output.

### Optimal Differentiation Condition

In order to identify the optimal differentiation condition, six differentiation medium compositions were tested and statistically analyzed (Figs. 1A-G). For all conditions, cells were plated at the same density on glass substrates and left to reach confluence in DMEM containing 10% FBS for three days to ensure comparability. After this point of time, which was defined as *t* = 0, the conditions were treated differently as described in Material and Methods. Briefly, the medium was changed to 1% FBS medium with or without regular addition of RA (*c*_RA_ = 10 − 1000 nM). Additionally, two conditions left with either 10% FBS and 1% FBS without further medium exchange were tested. Neither of those two has been investigated before, to the best of our knowledge.

**Figure 1.**
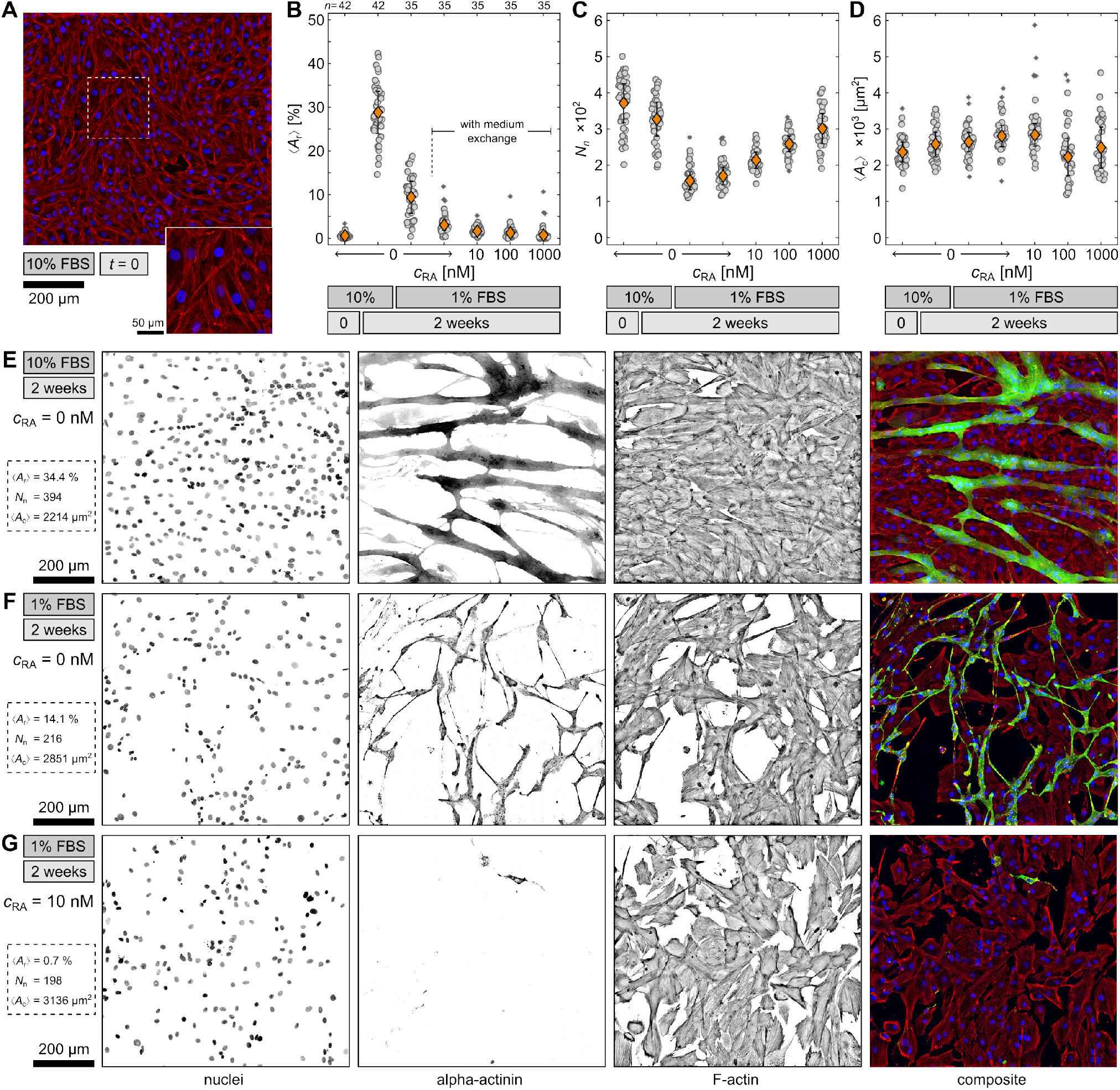
Comparison of differentiation conditions. (**A**) Example of undifferentiated H9c2 cells at *t* = 0. Shown are the nuclei (blue), sarcomeric alpha actinin (green), and F-Actin (red). (**B-D**) Quantification of differentiated cells by ratio between the area of differentiated cells ⟨*A*_r_⟩, the total number of nuclei *N*_n_, and the average differentiated cell area one nucleus occupies ⟨*A*_c_⟩ under six differentiation conditions. The orange diamonds represent the means with standard errors and the gray circles the data from individual 5 × 5 tiled images. The asterisks indicate outliers. Below each graph, the individual retinoic acid concentrations *c*_RA_, the respective FBS concentration and differentiation time are displayed. (**E-G**) Exemplary fluorescence stainings of differentiated H9c2 cells after 2 weeks at different conditions. Displayed are the nuclei, sarcomeric alpha actinin, F-actin, and the composite of all three channels.

For the comparison of the conditions, cells were fluorescence stained and observed on large 5 × 5 tiled images by confocal microscopy. From those images the differentiation rate was statistically quantified by an automated analysis routine (see Material and Methods). Figure 1A displays fluorescence-stained control cells at *t* = 0. As expected, the differentiation marker sarcomeric alpha actinin (αA) was not detected, and thus, the cells are still undifferentiated at this point. This can be confirmed by the area ratio ⟨*A*_r_⟩ between the area of differentiated cells (αA signal) and the area of all cells (F-actin signal), shown in Fig. 1B. For the undifferentiated cells, ⟨*A*_r_⟩ is close to 0%. Generally, ⟨*A*_r_⟩ represents the differentiation efficiency in terms of sarcomeric alpha actinin production. In this study, the highest ⟨*A*_r_⟩ of about 30% was found at the 10% FBS condition after two weeks, although previous studies indicate that serum reduction is necessary for differentiation of H9c2 (Ménard et al., 1999; Pereira et al., 2011; Suhaeri et al., 2015; Campero-Basaldua et al., 2023). However, the finding implies that FBS contains some components that support cell differentiation, and therefore, the differentiation rate is higher if the FBS concentration remains at 10%. In comparison, the 1% FBS condition without medium exchange exhibits only an ⟨*A*_r_⟩ of about 10%. Cells under this condition still form sarcomeric alpha actinin but less efficiently. In comparison, the 1% FBS condition with medium exchange leads to a significantly lower differentiation rate, despite the only difference being the frequency of medium exchange. A regular medium exchange seems to disturb cell differentiation as it possibly hinders cell-cell communication, which is necessary for effective differentiation. Consequently, all three conditions with RA display a very low ⟨*A*_r_⟩ as well, but also a relation between the respective RA concentration and ⟨*A*_r_⟩. The higher the RA concentration, the lower ⟨*A*_r_⟩ and hence the differentiation rate. In contrast to previous findings, the addition of RA seems to disturb or even inhibit the differentiation of H9c2, at least in terms of αA formation.

Nevertheless, an opposite trend for the RA conditions is observed for the total number of nuclei *N*_n_ (Fig. 1C). The higher *c*_RA_, the higher *N*_n_. While RA fails to stimulate sarcomeric alpha actinin production, it seems to promote proliferation instead. A similar impression can be gained when considering ⟨*A*_c_⟩, calculated as the total cell area divided by the total amount of nuclei (Fig.1D). While ⟨*A*_c_⟩ remains comparable to undifferentiated cells throughout all differentiation conditions, the two higher RA conditions exhibit a broader variation in ⟨*A*_c_⟩. Cells under these two conditions might be in various stages of the cell cycle, resulting in both small and normal-sized cells, which again suggests an increased proliferation activity. On the other hand, *N*_n_ is lowered for all reduced FBS conditions, indicating cell detachment, which could be explained by the lower amount of adhesion factors provided. In contrast, for the 10% FBS condition, *N*_n_ remains comparable to the nuclei number of undifferentiated cells at *t* = 0, implying dominant differentiation and cell fusion, as indicated by ⟨*A*_r_⟩ (Fig. 1B).

To illustrate the above-mentioned results further, Figs. 1E-G feature three typical examples of fluorescence-stained H9c2 cells after 2 weeks under the 10% FBS, 1% FBS, and 1% FBS with 10 nM RA conditions. The nuclei and F-actin channels demonstrate the difference in *N*_n_ and the total cell number respectively, whereas the alpha actinin channel exemplifies ⟨*A*_r_⟩. When comparing the cell morphology of the 10% FBS and 1% FBS condition without medium exchange, the cells exhibit an elongated shape and multiple nuclei in both. However, the 10% FBS condition leads to thick, voluminous cells, whereas the 1% FBS condition results in rather thin, spindle-like cells. This difference can also be detected in phase-contrast images (not shown). Since both conditions lack the addition of RA, the differentiated cells should be skeletal muscle cells according to the literature (Ménard et al., 1999; Branco et al., 2015). To confirm this assumption, these two conditions were investigated in more detail.

### Phenotype Determination

As indicated above, the influence of both differentiation conditions, i.e. 10% and 1% FBS, without RA and medium exchange was examined to determine whether the cells are cardiac or skeletal myocytes. In addition to αA, the cells were fluorescence stained with a cardiac Troponin T (cTT) antibody to distinguish heart and skeletal muscle cells. The cTT is highly specific for cardiac cells and only reacts with differentiated myocytes but not with undifferentiated myoblasts. Therefore, it also serves as an additional differentiation marker. Figure 2A compares ⟨*A*_r_⟩ in relation to both αA and cTT for control cells at *t* = 0 and differentiated cells after two weeks under the 10% and 1% FBS conditions without medium exchange. For both differentiation conditions, the cells are positive for cTT, and therefore, they are indeed cardiac muscle cells, contrarily to the initial assumption. Similarly, Fig. 2B illustrates the nuclei ratio *N*_r_, which is defined as the nuclei number of the differentiated cells divided by *N*_n_ and presents another independent way to quantify the differentiation efficiency. Both ⟨*A*_r_⟩ and *N*_r_ show comparable differentiation rates for αA and cTT within the same differentiation condition. Thus, all of the differentiated cells are cardiomyocytes; there is no indication of skeletal muscle cells, which we also exemplarily confirmed for the *c*_RA_ = 10 nM condition (see Fig. S1, supplemental material). This is in contrast to previous findings which claim that addition of RA is necessary to form heart cells (Ménard et al., 1999). Comparing the differentiation rate, *N*_r_ is even higher than ⟨*A*_r_⟩. Around 50% of the nuclei under the 10% FBS condition and 25% under the 1% FBS condition belong to differentiated cells, respectively. During differentiation, cells fuse and consequently possess multiple nuclei, however, the area does not correspond linearly to the number of nuclei per cell. For the control cells (*t* = 0), a slight difference between αA and cTT can be noticed for both ⟨*A*_r_⟩ and *N*_r_. Very few of the undifferentiated cells already expressed some level of cTT. On the one hand, cTT might be produced earlier than αA as soon as the cells reach confluence. On the other hand, a minority of H9c2 cells could occasionally produce cTT during some phases of the cell cycle and lose it again. To examine this hypothesis, undifferentiated myoblasts were observed one hour after cell seeding (see Fig. S2, supplemental material). At this point, no cardiac Troponin T was found. Hence, it is more likely that cTT is formed as soon as the cells reach confluence.

**Figure 2.**
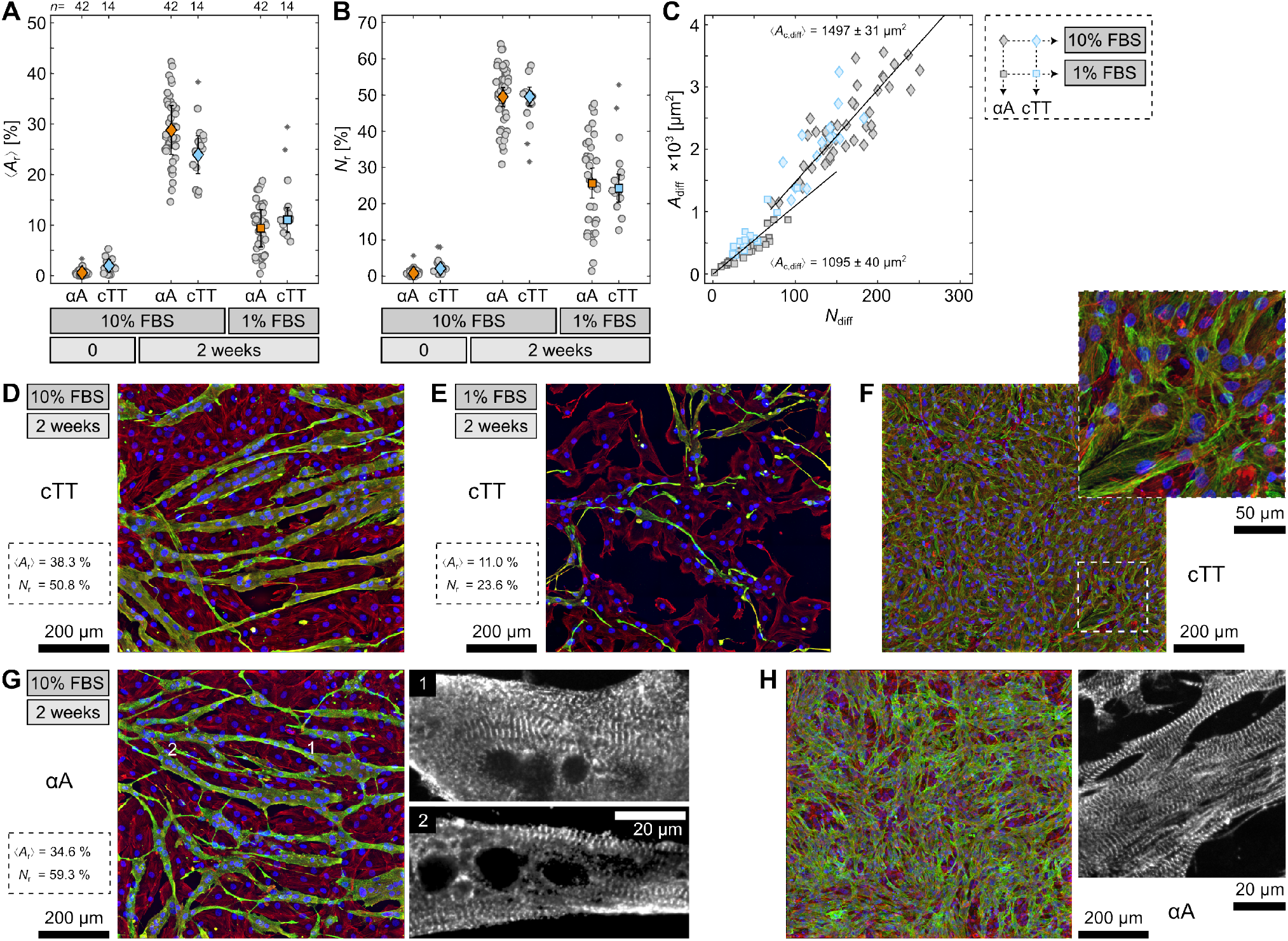
Quantification of sarcomeric alpha actinin (αA) and cardiac Troponin T (cTT) in differentiated H9c2 cells. (**A-B**) Comparison of the area ⟨*A*_r_⟩ and nuclei *N*_r_ ratio of differentiated cells for αA (orange) and cTT (blue) at different differentiation conditions. (**C**) Differentiated cell area *A*_diff_ versus number *N*_diff_ for αA (gray) and cTT (blue) at the 10% FBS (diamonds) and 1% FBS (squares) conditions. The slope of the solid lines (least squared fitted) indicates the average size differentiated cells occupy for each nucleus and is stated as ⟨*A*_c,diff_⟩. (**D, E, G**) Fluorescence stainings of differentiated H9c2 cells at 10% FBS and 1% FBS. (**F, H**) Fluorescence stainings of primary heart cells. Shown are the nuclei (blue), F-actin (red) and either cTT or αA (green), as indicated.

Additionally, differentiation with 10% FBS not only increases the differentiation rate as quantified by ⟨*A*_r_⟩ and *N*_r_ (Figs. 2A-B) but also leads to larger cells. Figure 2C displays a connection between the number of nuclei (*N*_diff_) and the area (*A*_diff_) of differentiated cells within one recorded image. The slopes indicate the average differentiated cell area one nucleus occupies (*A*_c,diff_) as about 1500 µm^2^ and 1100 µm^2^ for differentiation media with 10% and 1% FBS. Consequently, cells differentiated with 10% FBS are almost 50% larger than cells differentiated with 1% FBS. This confirms the impression already exemplified in Figs. 1E and 1F for αA. For cTT, the same difference in area size and morphology between the two conditions is depicted in Figs. 2D and 2E. Again, the 10% FBS cells (Fig. 2D) appear more voluminous and larger than the 1% FBS cells (Fig. 2E). Hence, if the FBS concentration is reduced during differentiation, the cells visibly exhibit signs of starvation. This starvation might result in stress and thus negatively influence the differentiation efficiency. For comparison, Fig. 2F shows cTT-stained primary cardiomyocytes which display comparable sarcomeric structures.

Another relation to primary cells appears when examining the differentiated 10% FBS cells more closely. Some cells developed sarcomere-characteristic striation (Fig. 2G). While the striation is hard to detect in the orthogonally projected images, it can be seen clearer when going through the individual focal heights (see enlargements in Fig. 2G). This finding strongly contradicts preceding studies which could not identify striation and even implied that differentiated H9c2 cells lack striation altogether (Suhaeri et al., 2015). In contrast to those studies, we used a differentiation condition that has not been considered before but led to a significantly improved differentiation. Comparing the 10% FBS cells with primary cardiomyocytes as regards αA again reveals strong likeness between the two (Fig. 2H). This even implies that differentiated H9c2 cells might be functioning cardiomyocytes.

### Cardiac Cell Dynamics

Due to the similarity of differentiated H9c2 cells to primary cardiomyocytes, the 10% and 1% FBS cells were examined for their calcium response, since it is part of the regulation of various functions such as contractility, hypertrophy, and gene expression (Aronsen et al., 2016). The differentiated cells were stained with the fluorescent dye Fluo-8 and stimulated by an electric field of about 10 V/cm to investigate the free intracellular Ca^2+^ concentration (Fig. 3A). The monophasic stimulation applied by an stimulation duration *τ*_s_ of 10 ms or 50 ms and a pacing period *T* ranging between 1000 ms and 240 ms. The recorded calcium signal was then used to extract the restitution properties of the cells at a single location. Figure 3B shows an example of the relation between calcium transient duration (CTD) and the calcium transient interval (CTI) for cells at 10% FBS after 10 days of differentiation. At 240 ms, the pacing was stopped since the signal-to-noise ration reached a critical level. This typical restitution property can also be illustrated using a signal oversampling analysis of the calcium transient, where individual waves are stacked and averaged (Hörning et al., 2017; Erhardt et al., 2024). Figure 3C displays three examples at different pacing periods. While the red line indicates the respective means, the black lines show the individual calcium transient curves. Even though the samples feature typical restitution properties for individual cells, i.e., a correlation between CTI and CTD for smaller *T*, and a CTD plateau for larger T, no wave conduction between the cells could be observed.

**Figure 3.**
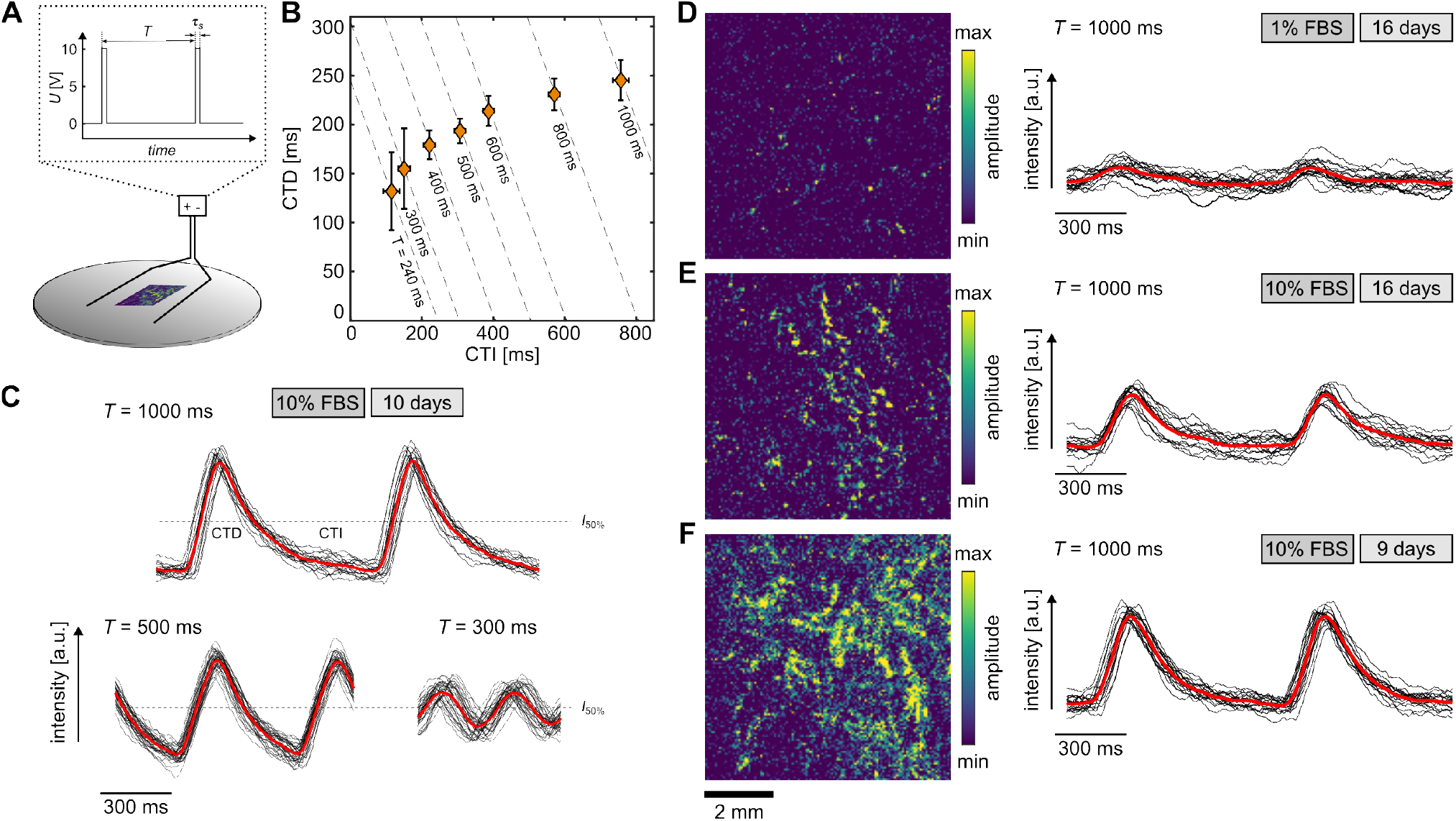
Calcium transients of differentiated H9c2 cells. (**A**) Scheme of experimental setup. The electric field of 10 V/cm is applied with a monophasic stimulation period *τ*_s_ of either 10 ms or 50 ms. (**B**) Restitution curve of differentiated H9c2 cells at 10% FBS after 10 days at *τ*_s_ = 10 ms. The data points show the mean values and the error bars the standard deviation. (**C**) Example of normalized calcium transience (red lines) for *T* = 1000 ms, *T* = 500 ms, and *T* = 300 ms for cells in (**B**). The black lines represent the periodically stacked individual calcium transients. (**D-F**) Examples of calcium transiences observed under different conditions and times. The left panels illustrate the amplitudes obtained by Fourier transformation imaging (FFI), and the right panels show the calcium transients of a single location. The examples are stimulated with *τ*_s_ = 50 ms and *T* = 1000 ms. The black lines represent the periodically stacked individual calcium transience and the red lines the normalized calcium transience.

For the 1% FBS condition, no wave conduction was detected either, and the calcium transients exhibited weaker signals, i.e., lower signal-to-noise ratio. Figure 3D illustrates such an example along with a normalized amplitude of a 6 × 6 mm^2^ field of view analyzed by Fourier transformation imaging (FFI) (Hörning et al., 2017). Calcium transients were observed only at very few locations in the tissue (yellowish color). In contrast, for the same age of the tissue (16 days) but the 10% FBS condition, active areas with stronger signals were identified (Fig. 3E). The strongest signal was detected after 9-10 days of differentiation (Figs. 3C and 3F), implying a maximum maturity before 2 weeks. Thus, the starvation over the long time period causes a reduction of active, functional cells. Nevertheless, the differentiated H9c2 cells under the 10% and 1% FBS conditions show calcium transients similar to primary heart cells and are therefore functional cardiac cells, at least in terms of their calcium response.

### Optimal Extracellular Matrix Condition

As glass does not represent the optimal extracellular matrix (ECM) condition for muscle cells, artificial ECMs, such as hydrogels, were used in the past. They better mimic natural ECM conditions(Hörning et al., 2012; Engler et al., 2006, 2004; Brock et al., 2022) and influence differentiation (Engler et al., 2004; Smith et al., 2018; Denes et al., 2019). In order to identify the optimal ECM condition for H9c2 cells, the response of single cells to polyacrylamide hydrogels with an E-modulus of *E* = 12.4 kPa were tested initially. This was chosen since it matches the natural rigidity of muscle cells (Engler et al., 2004). In a previous study, we established ECM conditions for the C2C12 myoblast cell line and observed the largest cell area at *ρ*_FN_ = 2.6 µg/cm^2^ on 12 kPa hydrogels (Brock et al., 2022). For the experiments, the cells were seeded in low densities on either plane or mechanically-roughened hydrogels, which were functionalized with different fibronectin densities (*ρ*_FN_ = 0.4 − 4.0 µg/cm^2^). After 24h, the cells were fluorescence stained and statistically analyzed. Figure 4A compares fluorescence-stained cells on plane hydrogels that were coated with 0.4 and 2.6 µg/cm^2^ fibronectin, respectively. The cells on the hydrogels with the lower fibronectin density appear smaller in size and more circular in shape. In contrast, the cells on the higher fibronectin density display a more complex spreading conformation by showing longer actin filaments, which indicates stronger adhesion to the coated substrate.

**Figure 4.**
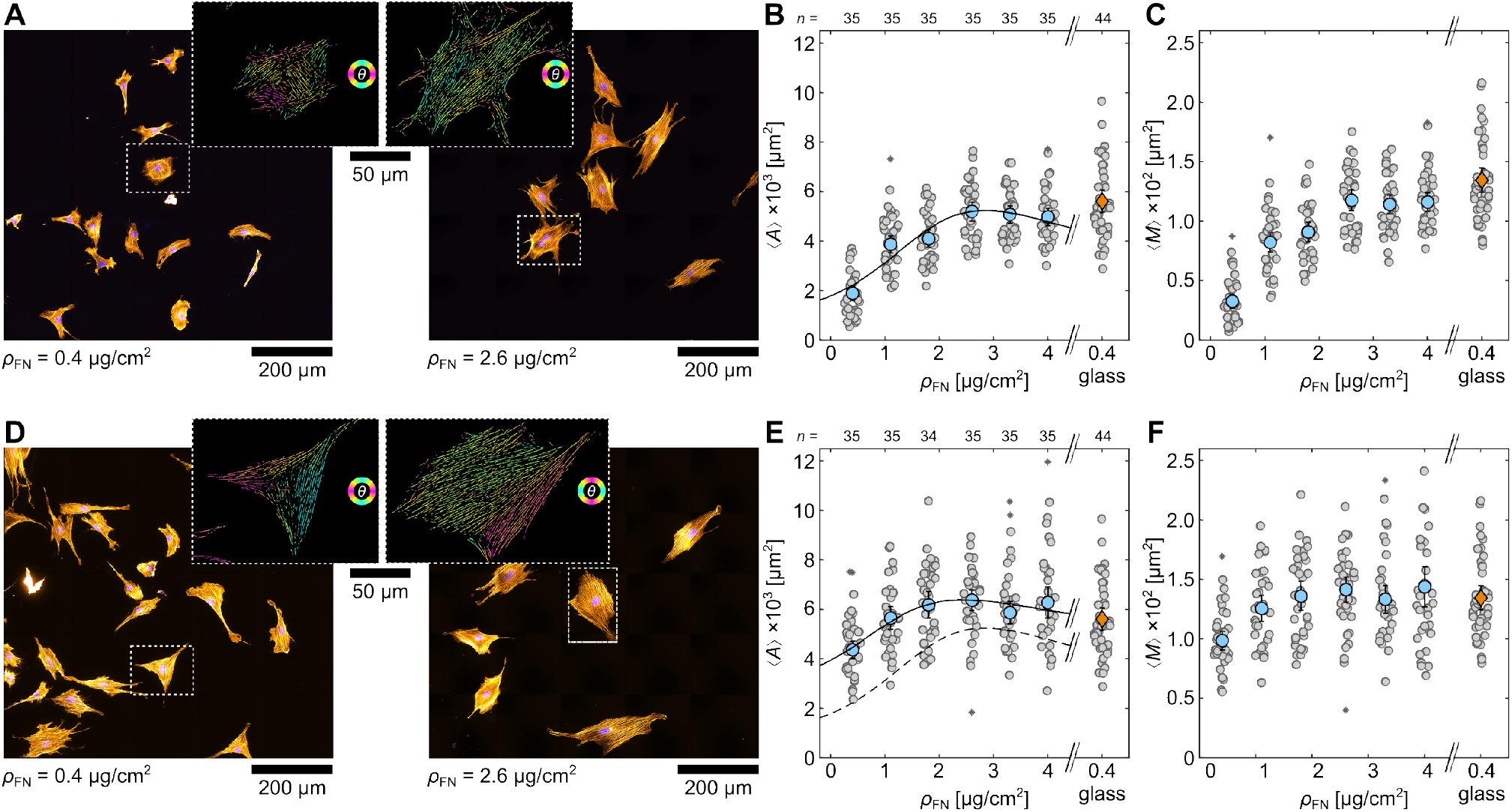
Single H9c2 cells on soft hydrogels coated with different fibronectin densities (*ρ*_FN_). Comparison between plane (**A**-**C**) and roughened hydrogel surfaces (**D**-**F**). (**A** and **D**) Fluorescence-stained cells on hydrogels (*E* = 12.4 kPa) with *ρ*_FN_ = 0.4 µg/cm^2^ (left) and *ρ*_FN_ = 2.6 µg/cm^2^ (right). The close ups illustrate two exemplary cells (dashed square) that are processed by the actin filament analysis. (**B-C** and **E-F**) Statistical analysis of mean projected cell area ⟨*A*⟩ and mean amount of actin per cell ⟨*M*⟩ on plane hydrogels for different *ρ*_FN_. The round gray data points are calculated from the individual 5 × 5 tiled images, and the orange diamond (glass) and blue circles (hydrogel) represent the mean with standard errors. The asterisks depict outliers.

To compare the morphological cell response, the mean projected cell area ⟨*A*⟩ and the mean F-actin amount per cell ⟨*M*⟩ for *ρ*_FN_ = 0.4 − 4.0 µg/cm^2^ was statistically quantified (Figs. 4B and 4C). Both ⟨*A*⟩ and ⟨*M*⟩ increase until about *ρ*_FN_ = 2.6 µg/cm^2^ and show a saturation thereafter. ⟨*A*⟩ can be parameterized by a Lorentz-like function, as

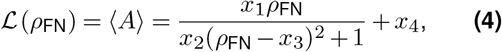

where *x*_*i*_ are fitting parameters. The black solid line represents the respective fit with an optimum at about *ρ*_FN_ = 2.6 µg/cm^2^ (Fig. 4B). This analysis has been applied before for C2C12 cells on collagen (Zemel et al., 2010) and fibronectin coated hydrogels (Brock et al., 2022). In the latter study, the maximum was also determined at *ρ*_FN_ = 2.6 µg/cm^2^. For comparison, ⟨*A*⟩ and ⟨*M*⟩ are also depicted for single H9c2 cells on glass, displaying similar values as cells on hydrogels with higher fibronectin density.

Since plane substrates do not promote the formation of confluent tissue, hydrogels (*E* = 12.4 kPa) that were mechanically roughened and thus feature micro scratches on their surface to support cell adhesion were tested, as a further step. Figure 4D illustrates examples of fluorescence-stained cells on roughened hydrogels at *ρ*_FN_ = 0.4 and 2.6 µg/cm^2^. Similar to cells on plane substrates, the cells on the lower fibronectin density seem slightly smaller than on the higher density. Nevertheless, the cells appear visibly larger and their actin filaments more oriented than on the plane hydrogels, especially for *ρ*_FN_ = 0.4 µg/cm^2^. The quantification of ⟨*A*⟩ and ⟨*M*⟩ for cells on roughened hydrogels (Figs. 4E and 4F) displays a similar trend as for plane hydrogels (Figs. 4B and 4C). ⟨*A*⟩ and ⟨*M*⟩ increase only up to *ρ*_FN_ = 2.6, which again indicates the optimum. Generally, roughened hydrogels lead to larger cells for all tested *ρ*_FN_, as indicated by the solid and dashed line in Fig. 4E. This difference is especially noticeable for the cells cultured on low *ρ*_FN_. At *ρ*_FN_ = 0.4 µg/cm^2^, for instance, the cells are about twice as large on the roughened hydrogels. Thus, the micro pattern influences cell adhesion. In conclusion, a more natural ECM, provided through the hydrogel, seems to be preferable for cells. As the mechanical roughening of the 12.4 kPa hydrogels in combination with a coating of *ρ*_FN_ = 2.6 µg/cm^2^ leads to a maximum in both ⟨*A*⟩ and ⟨*M*⟩ for single H9c2 cells, this ECM condition was chosen for subsequent experiments on cell differentiation.

### Differentiation on Soft Hydrogels

For the investigation of the differentiation dynamics of H9c2 cells on hydrogels, the optimum ECM condition, i.e., roughened hydrogel with *E* = 12.4 kPa and *ρ*_FN_ = 2.6 µg/cm^2^, in combination with the optimum medium condition (DMEM with 10% FBS) was used. Figures 5A-C show time-related comparisons between differentiation cells on glass and hydrogels at *t* = 0 and after 1 and 2 weeks of differentiation. The quantification of cells after 1 week was added since the cells expressed the highest calcium signals after 9-10 days of differentiation (Fig. 3). The cells were stained with the differentiation marker αA. Again ⟨*A*_r_⟩, *N*_r_, and *N*_n_ were statistically quantified (see Figs. 5 A-C). In comparison to glass substrates (orange diamonds), significantly lower differentiation rates were observed after 1 and 2 weeks for cells cultured on the optimized hydrogels (blue circles) for ⟨*A*_r_⟩ and *N*_r_ (Figs. 5D and 5E). Despite the natural substrate condition, the differentiation efficiency did not improve. Moreover, no striation could be detected for cells on hydrogels. However, in a few cases a comparable differentiation ratio was observed, as on glass. As opposed to this, the cells on glass reach their maximum differentiation ratio of about 40% after 1 week. Nevertheless, no sarcomere-typical striation could be found for those cells either, suggesting that their differentiation is not entirely completed after all and that a longer differentiation period is advantageous.

**Figure 5.**
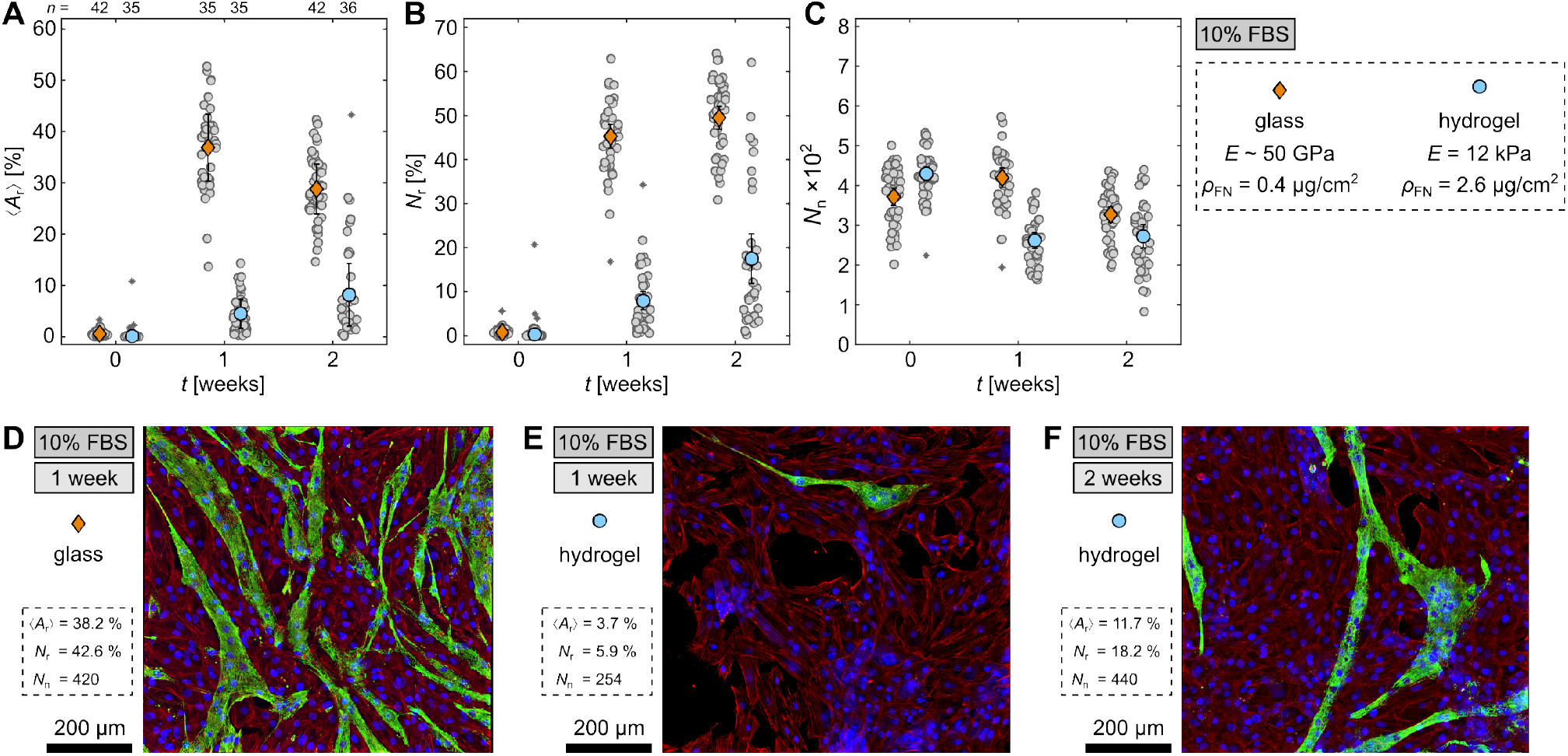
Differentiation of H9c2 cells on hydrogels. (**A-C**) Comparison of differentiated cells on glass and hydrogel with *E* = 12.4 kPa and *ρ*_FN_ = 2.6 µg/cm^2^ at 10% FBS. (**A-C**) Comparison of the area ⟨*A*_r_⟩ and nuclei *N*_r_ ratio of differentiated cells, and the total number of nuclei *N*_n_ per image. The gray circles represent the data from the individual 5 × 5 tiled images and the orange diamonds (glass) and blue circles (hydrogel) the means with standard errors. The asterisks indicate outliers. (**D-F**) Exemplary fluorescence-stained cells, depicting nuclei (blue), sarcomeric alpha actinin (green), and F-actin (red).

Regarding the total nuclei number *N*_n_ (Fig. 5C), the number of cells observed on both glass and hydrogel at *t* = 0 is comparable, suggesting sufficient adhesion to the substrate. During differentiation, however, the confluent cells on the hydrogels cluster and detach, resulting in lower *N*_n_ after both 1 and 2 weeks of differentiation. For primary cardiomyocytes, the roughened hydrogels provide sufficient adhesion to allow cardiac contraction (Erhardt et al., 2024), indicating that the proliferation and differentiation dynamics for H9c2 cells in confluent tissues are too strong, surpassing the adhesion forces and resulting in cell detachment. For comparison, *N*_n_ decreases on glass as well after 2 weeks, which indicates that some cells detach during differentiation independently of the substrate rigidity.

To visually illustrate the difference between the differentiation ratios on glass and hydrogels, Figs. 5D-F show fluorescence-stained cells after 1 week on glass (Fig. 5D) and hydrogel (Fig. 5E), as well as, after 2 weeks on hydrogel (Fig. 5F). There are fewer differentiated cells on both hydrogels compared to glass and also some holes between the cells on the hydrogels which matches the lower *N*_n_ and indicates detachment. Conclusively, the roughening of hydrogels enabled H9c2 cell differentiation, which was not possible on plane hydrogels, however, at a much lower differentiation efficiency than on glass substrates. Further improvements for cell attachment needs to be considered in future studies.

## Conclusions

Based on the statistical analysis, we showed that H9c2 cells differentiate most efficiently when left in 10% FBS medium for 1 week, whereas, sarcomeric striation was detected only after 2 weeks. Unlike other studies that propose serum reduction and addition of retinoic acid for successful differentiation, this study indicates the use of the 10% FBS medium composition to be advantageous. It is also easier to apply, reduces workload and is more cost-efficient since no medium change or supplements are necessary. Moreover, all cells that differentiated in only 10% and 1% FBS medium were positive for the cardiac Troponin T marker indicating differentiation to cardiac cells without the addition of RA, which again stands in contrast with results of previous studies (Ménard et al., 1999; Branco et al., 2015; Pereira et al., 2011). RA even seemed to inhibit cell differentiation, resulting in a significantly lower differentiation ratio for the three tested RA concentrations. This matches the results of another study with human myoblast cells that found an inhibition of RA on the expression of muscle differentiation markers such as Troponin T or myogenin, as well as, an upkeep of myoblast cells in an undifferentiated state (El Hadad et al., 2017). Similarly, another study recently discovered that differentiation of H9c2 cells into the cardiac phenotype is rather influenced by differentiation time than by addition of RA (Campero-Basaldua et al., 2023). In this connection, it should be noted that the passage number is also relevant for successful differentiation. For passages larger than 15, we observed that fewer cells appear morphologically differentiated, i.e., no elongated form and fewer cells with multiple nuclei. This might account for some studies, e.g. Patten et al. (2017), being unsuccessful in differentiation.

While other studies could not detect striation in the myocytes or even claimed that the H9c2 cell line does not exhibit striation (Suhaeri et al., 2015), the formation of striated myocytes for the 10% FBS condition was demonstrated in this study. It seems that a high serum level influences differentiation positively and provides the cells with nutrients or other factors that are necessary for striated muscle formation. As we identified striation in the differentiated cells, we tested the calcium response of differentiated H9c2 cells and demonstrated that these cells exhibit calcium dynamics similar to primary heart cells. This strongly suggests that the differentiated cells are functional heart cells. However, more investigation is necessary on this feature. Further cardiac characteristics could be examined, for instance, the membrane potential and contractility. Nonetheless, this result supports previous studies that found functional similarities to cardiac cells, such as the expression of cardiac-typical L-type calcium channels (Ménard et al., 1999; Hescheler et al., 1991) and cardiac-like responses to stimuli that cause hypertrophy (Watkins et al., 2011)

Finally, the use of hydrogels, that were expected to provide the cells with a more natural extracellular matrix, did not improve cell differentiation. One reason might be that the proliferation and differentiation dynamics are stronger than the adhesion forces and consequently lead to cell clustering and detachment. Nevertheless, we demonstrated that not only single C2C12 myoblasts (Brock et al., 2022) but also single H9c2 myoblasts prefer a fibronectin density of 2.6 µg/cm^2^. For confluent, proliferating cells, however, plane and even surface roughened hydrogels are not sufficient to maintain differentiating cell cultures. Thus, more sophisticated cell platforms are necessary. Promising approaches are 3D printed and generated surfaces that provide stronger adhesion properties, as exemplified before (Erben et al., 2020; Mei et al., 2019; Shabankhah et al., 2024).

## Supporting information

Supplemental Images

## Data Availability Statement

The raw data supporting the conclusions of this article will be made available by the authors, without undue reservation.

## Author Contributions

JB and MH conceptualized and designed the research and wrote the manuscript. JB performed experiments and data acquisition. MH implemented the analysis routines and analyzed the data. All authors contributed to the article and approved the submitted version.

## Acknowledgements

We would like to thank Julia Erhardt, Theresa Kühn, and Nadine Oder for the insightful discussions. Our thanks also go to Joachim Spatz and Cornelia Miksch for providing access to the Atomic Force Microscope (AFM) at the Max Planck Institute for Intelligent Systems, Stuttgart, Germany. We are also grateful to Stephan Eisler and Melanie Noack for their support and assistance with microscopic imaging at the Stuttgart Research Center of Systems Biology (SRCSB), University of Stuttgart.

