## Supplemental Images for "Optimization of H9c2 differentiation leads to calcium-active and striated cardiac cells without addition of retinoic acid"

bioRχiv

### Supplementary Material

#### 1 SUPPLEMENTARY TABLES AND FIGURES

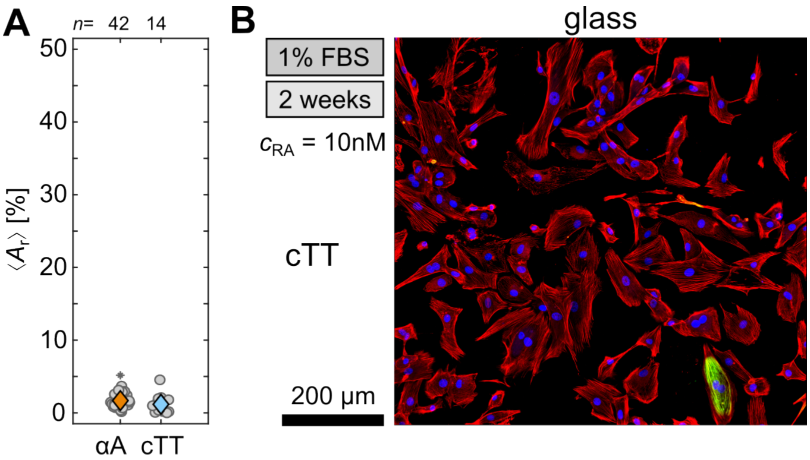


**Figure S1.** Comparison of sarcomeric alpha actinin (αA) and cardiac Troponin T (cTT) in differentiated H9c2 cells for the $c_{RA} = 1$ nM condition after 2 weeks. (**A**) Comparison of the ratio between the area of differentiated cells $\langle A_r \rangle$ for αA (orange) and cTT (blue). (**B**) H9c2 cells on glass observed 2 weeks after seeding on glass, depicting nuclei (blue), cTT (green), and F-actin (red).

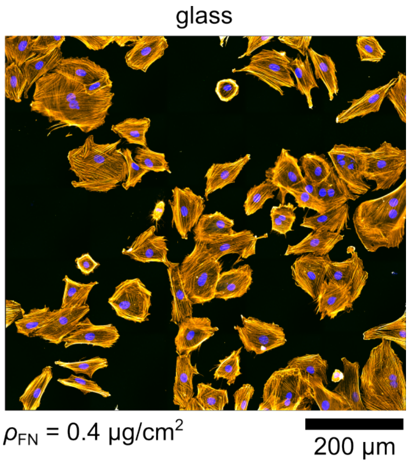


**Figure S2.** Undifferentiated H9c2 myoblasts observed one hour after seeding on glass, depicting nuclei (blue), cardiac Troponin T (green), and F-actin (orange). No cardiac Troponin T expressing cells have been detected.
